# Gut Microbiome Pattern Reflects Healthy Aging and Predicts Extended Survival in Humans

**DOI:** 10.1101/2020.02.26.966747

**Authors:** Tomasz Wilmanski, Christian Diener, Noa Rappaport, Sushmita Patwardhan, Jack Wiedrick, Jodi Lapidus, John C. Earls, Anat Zimmer, Gustavo Glusman, Max Robinson, James T. Yurkovich, Deborah M. Kado, Jane A. Cauley, Joseph Zmuda, Nancy E. Lane, Andrew T. Magis, Jennifer C. Lovejoy, Leroy Hood, Sean M. Gibbons, Eric S. Orwoll, Nathan Price

## Abstract

The gut microbiome has important effects on human health, yet its importance in human aging remains unclear. Using two independent cohorts comprising 4582 individuals across the adult lifespan we demonstrate that, starting in mid-to-late adulthood, gut microbiomes become increasingly unique with age. This uniqueness pattern is strongly associated with gut microbial amino acid derivatives circulating within the bloodstream, many of which have been previously identified as longevity biomarkers. At the latest stages of human life, two distinct patterns emerge wherein individuals in good health show continued microbial drift toward a unique compositional state, while the same drift is absent in individuals who perform worse on a number of validated health measures. The identified healthy aging pattern is characterized by an overall depletion of core genera found across most humans - primarily a depletion in the nearly ubiquitous genus *Bacteroides*. Consistently, retaining a high *Bacteroides* dominance into extreme age, or, equivalently, having a low gut microbiome uniqueness score, predicts decreased survival in a four-year follow-up. Our comprehensive analysis identifies the gut microbiome as a novel component of healthy aging, with important implications for the world’s growing older population.

## Introduction

The human gut harbors a diverse microbial ecosystem that has increasingly been shown to play an important role in host health^1–3^. Despite considerable progress in our understanding of the gut microbiome, very little is known about how it changes with age and how these changes interact with host physiology. Furthermore, there is no consensus on whether or not age-associated changes in the gut microbiome are related to the health state of the host. Importantly, identification of aging patterns within the gut microbiome could have major clinical implications for both monitoring and modifying gut microbiome health throughout life.

Several studies conducted on centenarian populations provided potential insight into gut microbial trajectories associated with aging. Biagi *et al*.^4^ demonstrated that gut microbiomes of centenarians (≤104 years of age) and supercentenarians (104+ years) show a depletion in core abundant taxa (*Bacteroides, Roseburia and Faecalibacterium*, among others), complemented by an increase in the prevalence of rare taxa. Similar findings have since been reported in other centenarian populations across the world, such as in Sardinian, Chinese and Korean centenarians, relative to healthy, younger controls^5–7^. Some studies have also reported higher α-diversity in centenarians compared to younger individuals^6–8^, suggesting that the gut microbiome continues to develop within its host even in the latest decades of human life.

Gut microbial associations reported in centenarians often contradict findings reported in younger elderly populations. In particular, studies on the ELDERMET cohort (i.e. the most extensively studied cohort of older persons with gut microbiome data to date) reported an increased dominance of the core genera *Bacteroides, Alistipes* and *Parabacteroides* in those 65+ years old compared to healthy, younger controls^9^. Studies on older long-term care residents further demonstrated a gradual change in microbiome composition associated with duration of stay in the care facility, which has been attributed to changes in diet and lifestyle^10,11^. Collectively, these and other studies^12,13^ provide a view of the human gut microbiome as relatively stable up until old age, at which point gradual compositional shifts occur that are driven by dietary and lifestyle changes, as well as declining health.

The often-contradictory findings in elderly and centenarian populations indicate there may exist multiple gut microbiome patterns of aging, some of which reflect better health and life expectancy outcomes than others. Although recent analyses have demonstrated a link between gut microbiome composition and long-term health outcomes ^3,14^, the scarcity of elderly cohorts with longitudinal follow-up data, the lack of detailed molecular phenotyping and health metrics, and the relatively small sample sizes of existing studies on aging limit our understanding of gut microbial changes seen across the human lifespan. In the present study, we overcome these limitations and present an analysis of the gut microbiome and phenotypic data from 4582 individuals spanning 18 to 98 years of age, with longitudinal follow-up data in an older cohort that allowed us to track survival outcomes.

## Results

We studied two distinct cohorts: a deeply phenotyped population of individuals who self-enrolled in a scientific wellness company (the ‘Arivale cohort’, ages 18-87) (**Table S1**) and the Osteoporotic Fractures in Men (MrOS) cohort (ages 78-98)^15–17^ (**Table S2**)(**Fig. 1**). These cohorts further subdivide into two groups each. The MrOS cohort separates into a discovery cohort (N=599) and a validation cohort (N=308), because stool samples from this population were processed in two separate batches several years apart. The Arivale cohort separates into Group A (N=2539) and Group B (N=1114), where the distinguishing factor is the use of different vendors for the collection and processing of stool samples (see Methods). We began by analyzing baseline data from the Arivale cohort to identify gut microbial aging patterns across most of the adult human lifespan, and investigate how these patterns correspond to host physiology. We then extended our analysis to the MrOS cohort, where we had detailed health metrics and follow-up data on mortality, to evaluate how the patterns identified within the Arivale cohort correspond to health and survival in the latter decades of human life.

**Fig. 1.**
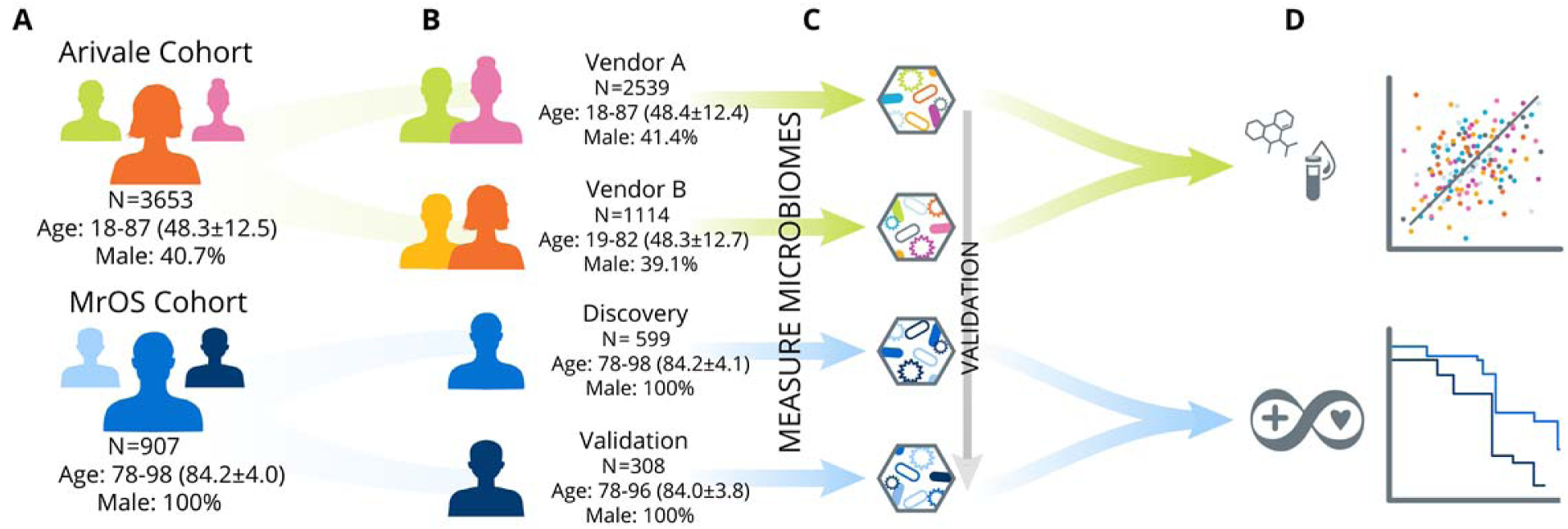
Conceptual outline of study and analysis workflow. **(A)** Two different study populations were used: the Arivale cohort and the Osteoporotic Fractures in Men (MrOS) cohort. **(B)** Each of these two study populations were further subdivided into two groups; the Arivale cohort was split based on the microbiome vendor used to collect and process samples while the MrOS cohort separated into Discovery and Validation groups based on the batch in which the samples were run (discovery samples were processed in the initial batch, validation samples were processed several years later). **(C)** We profiled the microbiomes from these four study populations beginning with the Arivale cohort and validating our findings across the three additional populations. (**D)** Our analysis pipeline further explored associations between the identified gut microbial aging pattern, lifestyle factors, and host physiology in the combined Arivale cohort, as well as health metrics and mortality in the combined MrOS cohort.

### A gut microbiome aging pattern that spans much of the adult lifespan

To characterize gut microbial patterns associated with aging, we initially performed a β-diversity analysis comparing all available baseline microbiome samples from a heterogenous, and relatively healthy Arivale population (**Fig**.**1** and **Table S1**). Our analysis involved extracting the minimum value for each individual from a calculated Bray-Curtis dissimilarity matrix. This value reflects how dissimilar an individual is from their nearest neighbor, given all other gut microbiome samples in the cohort. We refer to this as a measure of ‘uniqueness’: the higher the value, the more distinct the gut microbiome is from everyone else’s in the studied population. Arivale participants showed initial drift toward an increasingly unique gut microbiome composition starting between 40-50 years of age, and this continued to increase with every passing decade (linear models adjusted for age, body mass index (BMI), sex and Shannon diversity) (**Fig. 2A**). We replicated our analysis using additional β-diversity metrics. Uniqueness based on Weighted UniFrac demonstrated a similar positive association with age across both vendors, while Jaccard and Unweighted UniFrac metrics resulted in either a weaker association (vendor B) or no association (vendor A) with age (**Fig. 2B**). These results indicate that the observed age-related increase in uniqueness is likely not a result of the loss or acquisition of microbial genera in older individuals, which would increase unweighted β-diversity and Jaccard distance measures, but rather is driven more by shifts in relative abundance of microbes already present in the ecosystem.

**Fig. 2.**
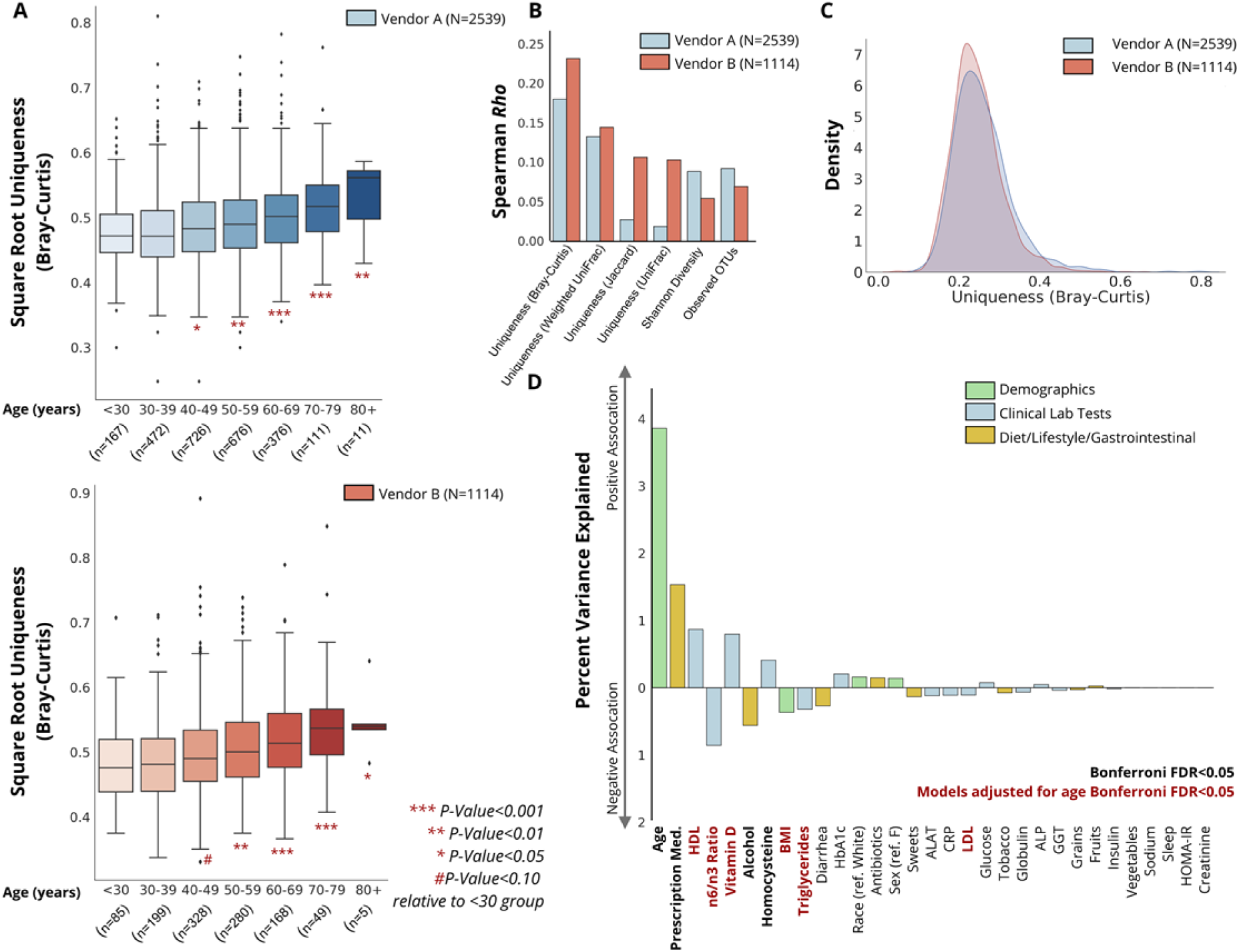
Associations between gut microbial uniqueness and age across the Arivale cohort. **(A)** Boxplots showing gut microbiome uniqueness scores calculated using the Bray-Curtis dissimilarity metric across the adult lifespan in individuals whose stool samples were collected and processed by vendor A (blue) or B (red). Asterisks indicate significant differences relative to the youngest <30 group, from a linear regression model adjusted for sex, BMI, and Shannon diversity. Box plots represent the interquartile range (25th to 75th percentile, IQR), with the middle line demarking the median; whiskers span 1.5 × IQR, points beyond this range are shown individually. (**B)** Spearman correlation coefficients for measures of uniqueness (β-diversity) and α-diversity with age in individuals whose stool samples were processed by vendor A or B. (**C)** Distribution of uniqueness calculated using the Bray-Curtis metric in each of the two vendors. **(D)** Percent of variance explained in Bray-Curtis uniqueness by a diverse number of demographic and lifestyle factors, as well as a subset of clinical laboratory tests.

To further characterize the observed gut microbiome aging pattern, and understand how it is reflected in host physiology and health, we combined data from both Arivale vendors (**Fig. 2C**) and tested the correspondence between Bray-Curtis uniqueness and a wide variety of clinical laboratory tests, demographic information, and self-reported lifestyle/health measures, adjusting for microbiome vendor (**Fig. 2D, Table S3**). Of all the factors tested, age demonstrated the strongest association with gut microbial uniqueness. Several other factors were significantly associated with uniqueness, but many of them were no longer significant after adjusting for age. In fact, after adjusting for age, essentially only lipid markers and BMI remained significantly associated with gut microbial uniqueness, with the direction of association indicating healthier metabolic and lipid profiles in individuals with more unique gut microbiomes: e.g. lower BMI, lower n6/n3 fatty acid ratio, higher high-density lipoprotein (HDL) cholesterol, lower low density lipoprotein (LDL) cholesterol, higher vitamin D, and lower triglycerides in individuals with more unique microbiomes (**Fig. 2D)**. Interestingly, self-reported dietary measures showed no association with our gut microbiome uniqueness score, suggesting that the identified gut microbial aging pattern is not driven by self-reported differences in dietary habits.

Women have an extended average lifespan compared to men ^18^, with previous studies also indicating varying aging patterns across sex ^19,20^. To evaluate whether the observed increased uniqueness with age is sex-dependent, we investigated the association of age with Bray-Curtis uniqueness independently in men and women, adjusting for age, BMI, Shannon diversity and microbiome vendor. Although both sexes showed a significant positive association between age and our gut microbiome uniqueness score, women showed a nearly 50% greater β-coefficient compared to men (adj. β (95%CI): men= 0.0088 (0.006, 0.012), women= 0.013 (0.010, 0.015), interaction term *P-Value*= 0.011), indicating that women’s microbiomes become more unique with age at a significantly faster rate.

### Reflection of gut microbial uniqueness in the host metabolome

Our research group has previously demonstrated a strong reflection of gut microbiome community structure in the human plasma metabolome^21^. In order to better understand how host physiology reflects the increasingly unique composition of the gut microbiome seen with aging, and to gain potential mechanistic insight into the functional changes that take place in the microbiota, we regressed our uniqueness measure against each of the 652 plasma metabolites measured in the Arivale cohort, adjusting for age, age squared, sex, a sex*age interaction term, BMI, vendor and Shannon diversity. A total of eight metabolites, all microbial in origin, remained significantly associated with uniqueness after multiple hypothesis correction (Bonferroni *P-Value*<0.05) (**Fig. 3A&B**). These metabolites fell primarily into one of two classes: phenylalanine/tyrosine metabolites (phenylacetylglutamine, p-cresol glucuronide, p-cresol sulfate) and tryptophan metabolites (3-indoxyl sulfate, 6-hydroxyindole sulfate and indoleacetate). Interestingly, significant changes in both tryptophan and phenylalanine pathways have been previously reported in centenarians relative to younger controls, with centenarians showing greater activation of these pathways in the gut microbiome ^22,23^. The previously identified longevity biomarker, phenylacetylglutamine ^23^, demonstrated the strongest correspondence with gut microbial uniqueness in our analysis, explaining 8.4% of the variance (adj. β (95%CI) = 0.015 (0.012,0.018), *P-Value*= 3.65e-20) (**Fig. 3C, Table S4**). These findings indicate that the observed gut microbial drift towards a more unique compositional state seen with age is characterized by alterations in microbial amino acid metabolism, which may serve as a useful biomarker for gut microbiome shifts across the human lifespan.

**Fig. 3.**
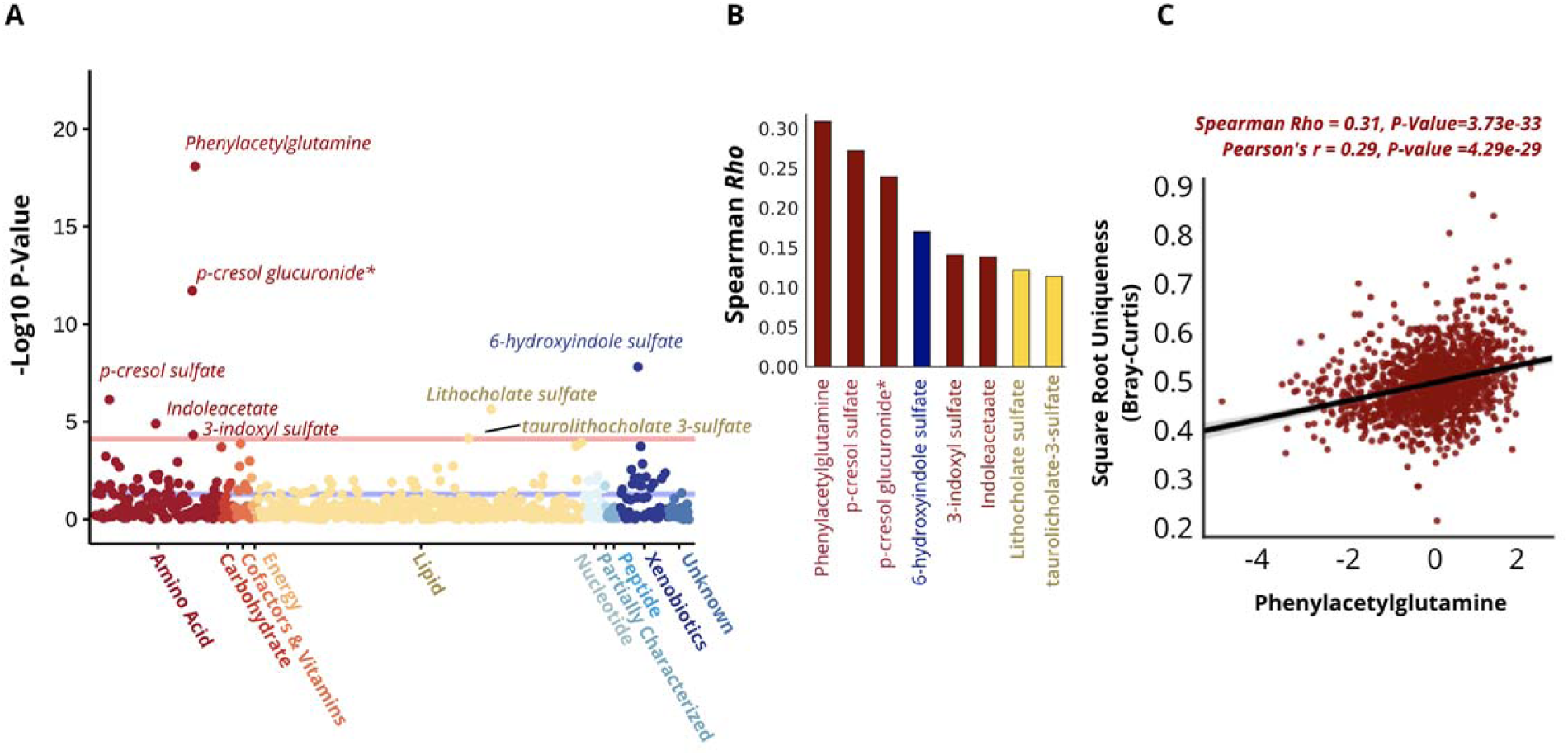
Reflection of gut microbiome uniqueness in plasma metabolites. **(A)** A plot of −log10 p-values for each of the 652 plasma metabolites measured in the Arivale cohort, from OLS regression models predicting Bray-Curtis uniqueness adjusted for age, age squared, sex, an age*sex interaction term, BMI, Shannon diversity and microbiome vendor. Metabolites are color-coded by their super-family. All metabolites above the red line are significant after multiple-hypothesis correction (Bonferroni P<0.05). * indicates metabolites that were confidently identified on the basis of mass spectrometry data, but for which no reference standards are available to verify the identity. **(B)** Spearman correlation coefficients for each of the metabolites significantly associated with Bray-Curtis uniqueness after adjusting for covariates and multiple-hypothesis correction (Bonferroni P<0.05). Bars are color-coded as in A). **(C)** Scatter plot of Bray-Curtis Uniqueness and the strongest metabolite predictor, phenylacetylglutamine. The line shown is a y∼x regression line, and the shaded regions are 95% confidence intervals for the slope of the line.

### Gut microbial pattern of healthy aging in latest decades of human life

To better understand the long-term health implications of the identified aging dynamics of the gut microbiome, we extended our analysis into a separate cohort of older men with paired health and longitudinal follow-up data (the MrOS cohort). The MrOS study recruited older male participants across the United States. At the fourth follow-up visit, a subset of the participants provided stool samples for 16S rRNA sequencing of their gut microbiome (discovery cohort N=599, validation cohort N=308)^17^. All participants who provided a stool sample exceeded 78 years of age at the time of sampling, allowing us to gain insight into the relationship between the gut microbiome and host health at the latest decades of human life (**Fig.1** & **Table S2**). Once again, we calculated a uniqueness score for each individual using the Bray-Curtis dissimilarity metric. Projecting MrOS microbiome data onto the first two Principal Coordinates revealed that samples with the highest Bray-Curtis uniqueness tended to fall away from common microbiome profiles, i.e. *Bacteroides* or *Prevotella* dominated ecosystems (**Fig. 4A-C**). In fact, the relative abundance of *Bacteroides* showed a strong negative association with gut microbiome uniqueness (Spearman Rho=0.73, **Fig. 4D**). The association was even more pronounced when the sum of both *Bacteroides* and *Prevotella* abundances for each individual was compared to gut microbiome uniqueness (Spearman Rho=0.80, **Figure S1A**).

**Fig. 4.**
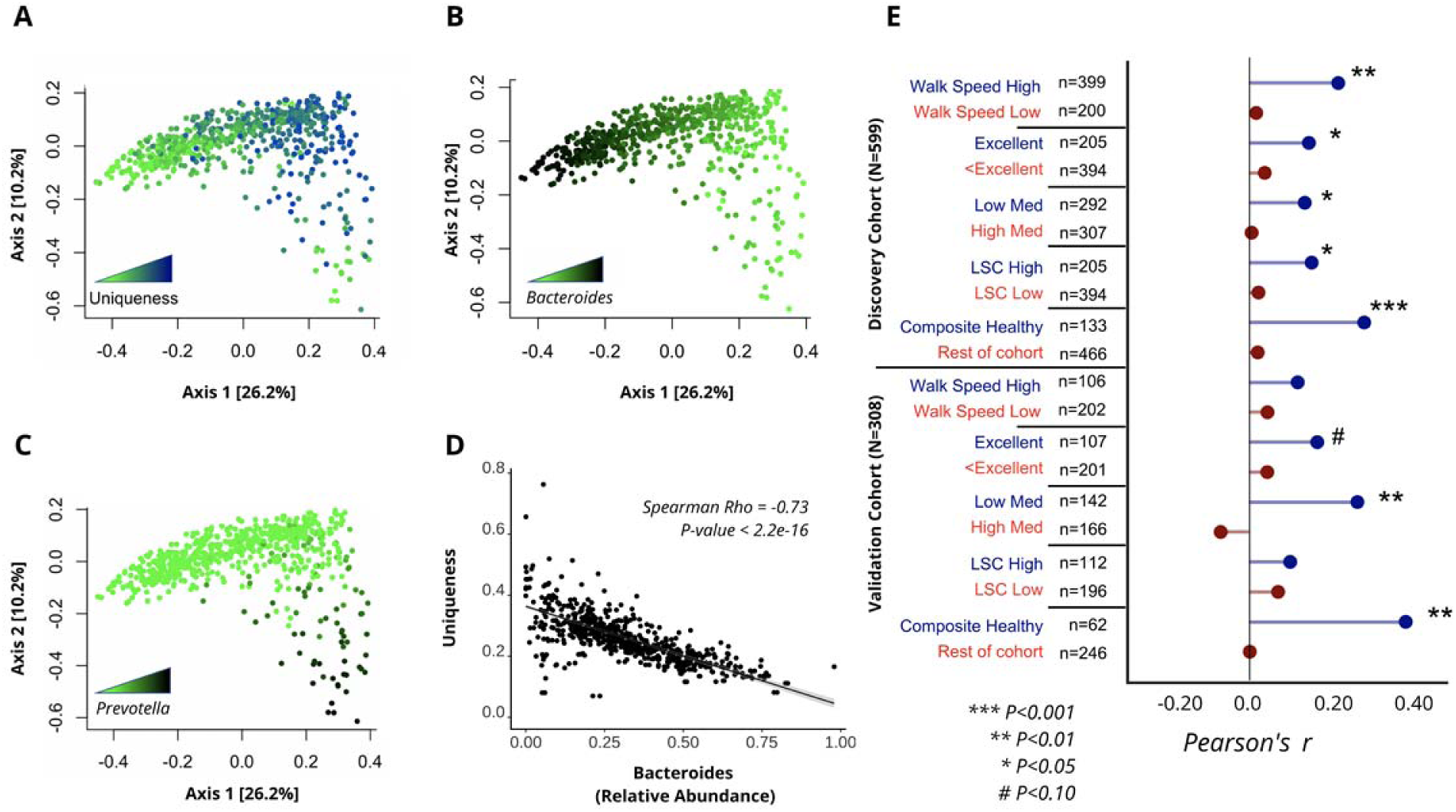
Increased dissimilarity of the gut microbiome as a function of healthy aging in the MrOS Cohort. **(A-C)** PCoA of the MrOS discovery cohort color-coded by **(A)** Bray-Curtis uniqueness, **(B)** relative *Bacteroides* abundance, and **(C)** relative *Prevotella* abundance. **(D)** Scatter plot demonstrating the negative association of the relative abundance of the most dominant genus *Bacteroides* and gut microbial uniqueness in the discovery cohort. The line shown is the y∼x regression line, while the shaded region corresponds to the 95% confidence intervals for the slope of the line. (**E)** Correlation of Bray-Curtis uniqueness scores with age across the MrOS discovery and validation cohorts under different health stratifications. ‘Excellent’ corresponds to individuals who self-reported their health to be excellent, while ‘<Excellent’ incorporates all individuals who self-reported their health being anything less than excellent (good, fair, poor, or very poor).’Composite Healthy’ refers to individuals who fell into the healthy sub-group in at least 3 of the 4 stratifications performed. LSC: Life-Space Score.

Consistent with our initial analysis, age showed a trending positive association with our uniqueness score in the MrOS cohort (Pearson’s r=0.075, *P-Value*= 0.065). Unlike the Arivale cohort, MrOS participants were considerably more health heterogenous at time of sampling, with a large proportion of participants reporting chronic conditions (**Table S2**). The large health heterogeneity of MrOS participants provided an opportunity to better understand whether the observed increase in microbiome dissimilarity with age depends on host health. Hence, we re-ran the above analysis under four different stratifications based on: medication use, self-perceived health, life-space score (LSC), and walking speed. We chose these four health metrics because collectively they encompass a diverse repertoire of health in older populations (**Table 1**).

**Table 1:**
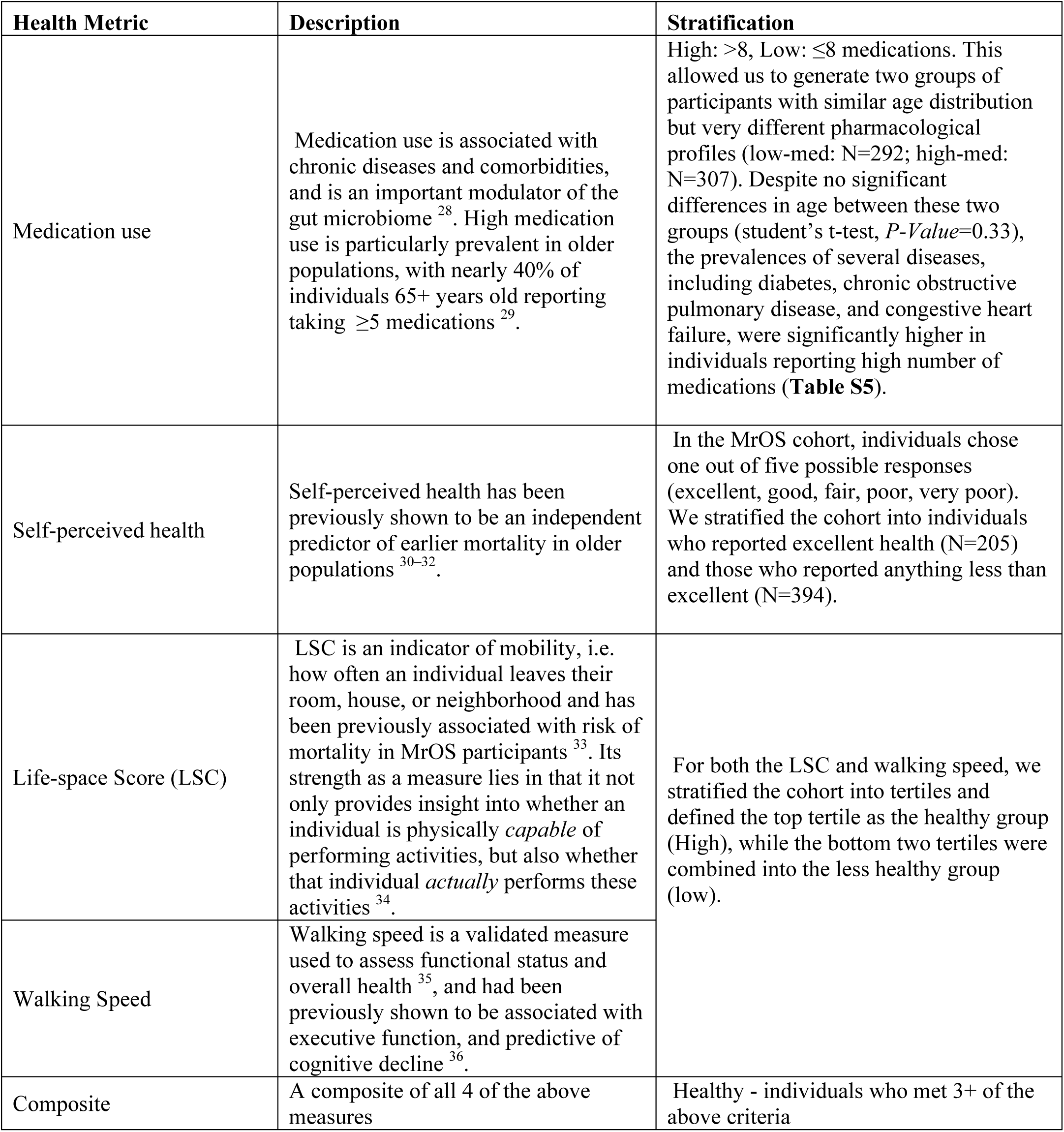
Description of health metrics used for stratification in the MrOS cohort.

| Health Metric | Description | Stratification |
| --- | --- | --- |
| Medication use | Medication use is associated with chronic diseases and comorbidities, and is an important modulator of the gut microbiome<sup>28</sup>. High medication use is particularly prevalent in older populations, with nearly 40% of individuals 65+ years old reporting taking $\geq 5$ medications<sup>29</sup>. | High: $>8$, Low: $\leq 8$ medications. This allowed us to generate two groups of participants with similar age distribution but very different pharmacological profiles (low-med: N=292; high-med: N=307). Despite no significant differences in age between these two groups (student's t-test, $P$-Value=0.33), the prevalences of several diseases, including diabetes, chronic obstructive pulmonary disease, and congestive heart failure, were significantly higher in individuals reporting high number of medications (**Table S5**). |
| Self-perceived health | Self-perceived health has been previously shown to be an independent predictor of earlier mortality in older populations<sup>30–32</sup>. | In the MrOS cohort, individuals chose one out of five possible responses (excellent, good, fair, poor, very poor). We stratified the cohort into individuals who reported excellent health (N=205) and those who reported anything less than excellent (N=394). |
| Life-space Score (LSC) | LSC is an indicator of mobility, i.e. how often an individual leaves their room, house, or neighborhood and has been previously associated with risk of mortality in MrOS participants<sup>33</sup>. Its strength as a measure lies in that it not only provides insight into whether an individual is physically *capable* of performing activities, but also whether that individual *actually* performs these activities<sup>34</sup>. | For both the LSC and walking speed, we stratified the cohort into tertiles and defined the top tertile as the healthy group (High), while the bottom two tertiles were combined into the less healthy group (low). |
| Walking Speed | Walking speed is a validated measure used to assess functional status and overall health<sup>35</sup>, and had been previously shown to be associated with executive function, and predictive of cognitive decline<sup>36</sup>. |  |
| Composite | A composite of all 4 of the above measures | Healthy - individuals who met 3+ of the above criteria |

Under all stratifications considered, we observed a stronger positive association between age and microbiome uniqueness in healthier individuals, while the association was absent altogether in individuals demonstrating worse health (**Fig. 4E**). We further generated a composite stratification (composite healthy), where MrOS participants had to meet at least three of the four criteria outlined above to be classified as healthy (**Table 1 & Table S2**). In this limited group of 133 individuals we observed an even stronger association between gut microbial uniqueness and age than under any individual stratification. We replicated the analysis on the second batch of MrOS gut microbiome samples, which were processed independently several years apart (validation cohort, N=308), demonstrating very similar results (**Fig. 4E**). We also ran the same analysis using Weighted UniFrac dissimilarity, and observed high level of congruence between results (**Fig. S1B**). In contrast, measures of α-diversity were not significantly associated with age under any stratification considered (**Fig. S1B**).

### Gut microbiome and mortality in extreme aging

Next, we focused exclusively on community-dwelling individuals (i.e. excluding participants in assisted living, nursing homes, and/or who have been hospitalized in the past 12 months) from the two MrOS data sets, combined together for increased power (N=706) (**Fig. 1**). We performed genus-level differential abundance analysis to identify genera associated with age in healthy composite individuals (N=173) and the remainder of the cohort (N=533), separately, adjusting for batch (discovery/validation) and BMI. In healthy composite individuals, only the genus *Bacteroides* (adj. β (s.e.): −0.062 (0.017), *P-Value*=0.0006) demonstrated a significant negative association with age after multiple hypothesis correction (**Fig. 5A**). These findings are consistent with our gut dissimilarity analysis, where the uniting feature of unique microbiomes is the depletion of the most common and dominant genera. Consistently, there was no significant association between age and *Bacteroides* in participants who did not meet our health criteria (adj. β (s.e.): −0.008 (0.009), *P-Value* =0.37) (**Fig. 5A**). In contrast, individuals in worse health demonstrated a distinct gut microbiome aging pattern characterized by a decline in the genera *Lachnoclostridium* (adj. β (s.e.): −0.035 (0.0091), *P-Value* =0.0002) and the *Rumminococace family* genus *UBA1819* (adj. β (s.e.): −0.074 (0.015), *P-Value* =2.57e-06) with age. These results provide further evidence for the existence of multiple gut microbiome aging patterns in the later stages of human life.

**Figure 5.**
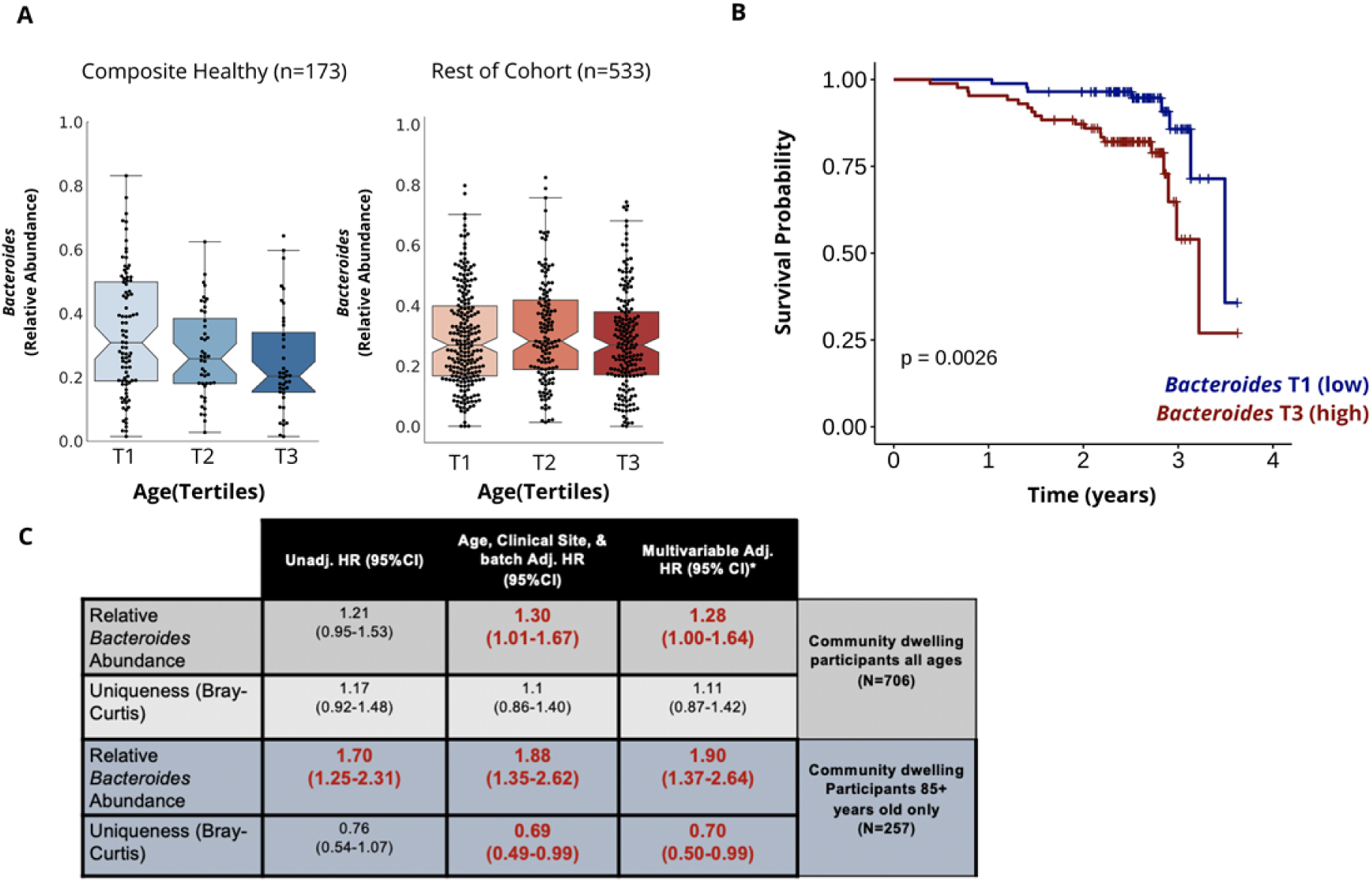
Association of *Bacteroides* abundance and survival in latter decades of human lifespan. **(A)** Boxplots demonstrating the relative abundance of the genus *Bacteroides* across tertiles of age in community-dwelling individuals identified as healthy on 3+ criteria specified (composite healthy) (n=173) and the remainder of the cohort (n=533). **(B)** Kaplan Meier Curve demonstrating the association between overall survival and relative *Bacteroides* abundance grouped into tertiles in community-dwelling MrOS participants who were 85+ years at time of sampling (N=257). **(C)** Unadjusted, age, clinical site and batch adjusted and multivariable adjusted Hazard Ratios (HR) of relative *Bacteroides* abundance and Bray-Curtis Uniqueness scores from Cox Proportional Hazard Regression models evaluating mortality risk in all community-dwelling MrOS participants and exclusively community-dwelling MrOS participants 85+ years old. Multivariable models were adjusted for age, clinical site, BMI, self-perceived health, diagnosis of congestive heart failure, and batch in which stool samples were processed. Both relative *Bacteroides* abundance and The Bray-Curtis uniqueness measures were scaled and centered prior to mortality analysis. Significant HRs are bolded and colored in red (*P*≤*0*.*05*).

Given that our findings from both β-diversity and differential abundance analysis of healthy elderly are consistent with observations previously reported in centenarians ^4^, we utilized longitudinal data from the MrOS cohort to investigate whether the observed gut microbiome pattern of healthy aging is predictive of mortality. We performed the analysis in two steps: 1) on all community-dwelling participants (N=706) and 2) only on community-dwelling participants in the top age tertile (85+ years of age, N=257) at time of gut microbiome sampling, because these participants were the closest to achieving extreme age in the course of the study’s follow-up period (∼4 years). When focusing on all individuals in the cohort, we identified a significant positive association between relative *Bacteroides* abundance and increased risk of all-cause mortality, independent of age, BMI, clinical site, self-perceived health, diagnosis of congestive heart failure, and batch in which stool samples were processed. Replicating the analysis in the oldest individuals (85+ years old) revealed a stronger association and higher Hazard Ratios compared to the whole cohort (**Fig. 5B-C**). Using the participants’ calculated Bray-Curtis uniqueness score yielded comparable results in 85+ year olds, where mortality risk decreased in individuals with more unique gut microbiomes independent of the same covariates. In contrast, the same association between Bray-Curtis uniqueness and mortality was not present when younger participants were included in the analysis (**Fig. 5C**).

## Discussion

There is a limited understanding of how the human gut microbiome changes throughout adulthood and how these changes influence host physiology. Here, we evaluated gut microbial patterns associated with aging across 4582 individuals from two distinct study populations spanning 18-98 years of age. The major findings of our analysis were: 1) individual gut microbiomes became increasingly more unique with age, starting in mid-to-late adulthood, and this uniqueness was positively associated with known microbial metabolic markers for health and longevity; 2) the increase in microbiome uniqueness with age occurred in both males and females, but was 50% more pronounced in females; 3) in the later decades of human lifespan, healthy individuals continued to show an increasingly unique gut microbial compositional state (associated with a decline in core taxa) with age, while that pattern was absent in those in worse health; 4) in individuals approaching extreme age (85+ years old), retaining high relative *Bacteroides* abundance and having a low gut microbiome uniqueness score were both associated with decreased survival in the course of 4 year follow-up. These observations are strengthened by the presence of similar age-related trends in two separate cohorts, and the replication of associations with health and longevity in a validation cohort.

Our findings indicate that healthy aging of the gut microbiome involves depletion of core microbes and their replacement by less common taxa, resulting in increasingly distinct microbiomes. These findings are consistent with patterns previously reported in centenarians across the world ^6,7^, despite the fact that dominant genera (i.e. core microbiota) often vary across cultures and geographic locations ^24^. Using our alternative beta-diversity approach, we provide novel insight into the aging gut microbiome that a) validates across different vendors and cohorts and b) is consistent with previous longevity research. It is quite possible that becoming increasingly dissimilar as you age is a universal characteristic, independent of the variability in core gut microbes observed across the world (e.g. *Bacteroides* vs. *Prevotella*). This would make gut microbiome uniqueness an intriguing new dimension of healthy aging, and a critical new component for personalized medicine and precision health.

The reflection of gut microbial uniqueness in plasma phenylalanine/tyrosine and tryptophan microbial metabolites is consistent with our recent work showing a robust relationship between the host blood metabolome and gut microbial diversity ^21^. Both tryptophan and phenylalanine metabolism have been implicated in longevity ^22,23,25^. Phenylacetylglutamine and p-cresol sulfate demonstrated some of the strongest associations with gut microbial uniqueness, independent of age and other covariates. These same metabolites were previously proposed as biomarkers for healthy aging and longevity ^23^. Additional metabolites associated with our observed gut microbial pattern were dominated by indoles, which are gut microbiome degradation products of tryptophan. Indoles have been shown to increase healthspan and extend survival in a number of animal models ^26^. Their most characterized mechanism of action is mediating inflammation through binding the aryl hydrocarbon receptor ^27^. While further studies are needed to establish a direct link between these microbial compounds and longevity in humans, the elevated levels of these metabolites in circulation in individuals whose microbiomes are more unique opens promising new leads into the role of the gut microbiome in aging.

Previous studies in older populations have suggested that gut microbial composition and structure is generally stable throughout adulthood and into old age ^12^, at which point changes are observed and further accelerated due to adverse health events and lifestyle changes (i.e. entering long-term care facilities) ^10,11,13^. In sharp contrast, our findings suggest that gut microbiomes of healthy individuals continue to develop throughout aging, and that it is the lack of this development that appears to be associated with worse health and prognosis. As our understanding of the aging microbiome increases, monitoring and identifying modifiable features that may promote healthy aging and longevity will have important clinical implications for the world’s growing older population.

## Acknowledgments

We thank C. Funk for helpful discussions throughout the course of this project. We also thank J. Dougherty and M. Brunkow for their coordination efforts.

## Author contributions

T.W., S.M.G., L.H., E.S.O., and N.P conceptualized the study. T.W., J.W., J.L, J.A.C., S.M.G., and E.S.O participated in study design. T.W., C.D., N.R., S.P., J.W., J.L., J.C.E., A.Z., and J.T.Y. performed data analysis and figure generation. G.G. and M.R. aided in dissimilarity analysis. G.G., M.R., N.E.L., J.Z., J.A.C and D.M.K. assisted in results interpretation. A.T.M. and J.L. managed the logistics of data collection and integration. T.W., S.M.G., E.S.O and N.P. were the primary writers of the paper, with contributions from all authors. All authors read and approved the final manuscript.

## Data and materials availability

Qualified researchers can access the full Arivale deidentified dataset analyzed in this study for research purposes. Requests should be sent to. The MrOS data set is available to researchers through the following website: https://mrosdata.sfcc-cpmc.net.

## Methods

### Cohorts

The Arivale cohort consists of individuals over 18 years of age who between 2015 and 2019 self-enrolled in a now closed scientific wellness company. The cohort has been described in detail previously ^15^. For this study, only baseline measurements were considered for each participant. The only inclusion criterion was the availability of gut microbiome data in order to maximize the number of gut microbiomes to which each sample is compared. Demographic information on the cohort is provided in Table S1.

The MrOS study is an ongoing prospective study of close to 6000 men recruited across six clinical U.S. sites. The cohort, recruitment criteria, and stool sample collections have been previously described in detail ^16,17^. Briefly, during the fourth follow-up visit of the original study, a subset of participants across all six clinics was asked if they would consent to have their stool sampled for microbiome analysis. Participants who agreed were given the OMNIgene-GUT stool/feces collection kit (OMR-200, DNA Genotek, Ottawa, Canada) and collected the fecal sample at their homes. Demographic information on MrOS participants is provided in Table S2. In the initial uniqueness analysis, all participants with available high-quality microbiome data were used for analysis (N=907). Subsequent differential abundance analysis focused exclusively on community-dwelling individuals (N=706) (excluding individuals in assisted living, nursing homes and who have been hospitalized in the past 12 months). Finally, survival analysis was conducted on all community dwelling individuals as well as specifically on community dwelling individuals in the latest stages of aging (85+ years old, N=257). The number of deaths in the whole community dwelling group and in 85+ year old community dwelling group was 66 and 41, respectively.

### Microbiome Analysis

#### Arivale cohort

Analysis of gut microbiome data from the Arivale cohort has been described in detail elsewhere ^21,37^. Briefly, independent of the vendor used, stool samples were collected at the participants’ homes using DNA collection kits with a proprietary chemical DNA stabilizer to maintain DNA integrity at ambient temperatures following collection. Gut microbiome sequencing data in the form of FASTQ files were provided on the basis of either the 300-bp paired-end MiSeq profiling of the 16S V3 + V4 region (DNAgenotek, vendor A) or 250-bp paired-end MiSeq profiling of the 16S V4 region (Second Genome, vendor B). Further analysis was performed using the denoise workflow from mbtools (https://github.com/gibbons-lab/mbtools) that wraps DADA2. In summary, we first trained DADA2 ^38^ error models separately for each sequencing run and used those to obtain sequence variants for each sample. This was followed by de novo chimera removal which removed ∼17% of all reads as chimeric and resulted in about 89,000 final sequence variants across all samples. Taxonomy assignment was performed using the RDP classifier with the SILVA database (version 132). Here 99% of the reads could be classified on the family level, 89% on the genus level and 32% on the species level. Species level taxonomy was identified by exact alignment to the SILVA reference sequences. Sequence variants were aligned to each other using DECIPHER ^39^ and the multiple sequence alignment was trimmed by removing each position that consisted of more than 50% gaps. The resulting core alignment had a length of 420 base pairs and was used to reconstruct a phylogenetic tree using FastTree ^40^. Downstream gut microbiome data analysis was conducted using the *Phyloseq* Package ^41^. In two separate analyses, gut microbiome samples were rarefied to 13703 (vendor A, DNAGenotek) and 39810 (Vendor B, Second Genome) reads, the minimum number of reads per sample for each vendor. For uniqueness analysis, the Bray-Curtis ^42^, Unweighted and Weighted UniFrac ^43^, and Jaccard matrices were calculated for all samples within each vendor using the rarefied Genus table. The minimum value for each row, corresponding to the dissimilarity of each sample to their nearest neighbor, was then extracted from the matrix and used for downstream analysis.

#### MrOS cohort

Stool samples were processed at the Alkek Center for Metagenomics and Microbiome Research (CMMR) at Baylor College of Medicine using their custom analytic pipeline in two separate batches (Discovery N=599, Validation N=320). 16Sv4 rDNA amplicon sequences were clustered into Operational Taxonomic Units (OTUs) at a similarity cutoff value of 97% using the UPARSE algorithm ^44^. OTUs were then mapped to an optimized version of the SILVA Database ^45^ containing only the 16S V4 region to determine taxonomies. Abundances were recovered by mapping the demultiplexed reads to the UPARSE OTUs ^44^. Preliminary microbiome data analysis was conducted using the *Phyloseq* Package. For α-diversity and uniqueness analysis, OTUs were rarefied to 9424 reads, which is the minimum number of OTUs/sample in the discovery cohort. The same rarefaction number (9424) was used in the Validation cohort (N=320). A total of 12 samples had less reads than the specified cut-off, and hence were excluded from the analysis (Validation N=308). α-diversity measures were calculated at the OTU level using the Phyloseq package ^41^. For ß-diversity analysis, OTUs were collapsed into genera. Uniqueness was calculated as described for the Arivale cohort. The calculated uniqueness measure for each participant was then used for downstream analysis. As part of our analytical pipeline, we also performed differential abundance analysis assessing the relationship of individual genera with age in individuals defined as healthy and unhealthy, separately. Analysis was performed in R (version 3.44) using beta-binomial regression through the Corncob package (version 1.0) ^46^. Models were adjusted for BMI, and batch (discovery/validation). Type 1 error was controlled using the Bonferroni method (P<0.1).

### Plasma Metabolomics & Clinical Laboratory Tests

Blood draws for all assays were performed by trained phlebotomists at LabCorp or Quest service centers. For the 24-hour period leading up to the blood draw, Arivale participants were required to avoid alcohol, vigorous exercise, aspartame and monosodium glutamate, and to begin fasting 12 hours in advance. Plasma metabolomics assays were conducted on the samples by Metabolon, Inc. Sample handling, quality control and data extraction, along with biochemical identification, data curation, quantification and data normalizations have been previously described ^37^. For analysis, the raw metabolomics data were median scaled within each batch, such that the median value for each metabolite was one. To adjust for possible batch effects, further normalization across batches was performed by dividing the median-scaled value of each metabolite by the corresponding average value for the same metabolite in quality control samples of the same batch. In this study, we analyzed participants’ baseline plasma metabolomics data. A 10% missing value threshold was set, which was passed by 652 metabolites. Missing values for metabolites were imputed to be the minimum observed value for that metabolite. A total of 1476 Arivale participants had paired gut microbiome-plasma metabolome data. Values for each metabolite were log transformed prior to analysis. Clinical laboratory tests were conducted by either Quest or LabCorp. A 10% missing value threshold was set for each clinical laboratory test used in the analysis. All but 104 participants (N=3549) had paired clinical laboratory-gut microbiome data. Both metabolomics and clinical laboratory tests were scaled and centered prior to analysis and only baseline measures for each individual were used.

### Lifestyle/Health Questionnaires in the Arivale Cohort

Data on lifestyle, diet and health were obtained through self-administered questionnaires completed by Arivale participants during their initial assessment. For reporting antibiotic use, participants chose from three possible responses (‘not in the past year’, ‘in the past year’ and ‘in the past three months’) which were recoded into ordinal variables 0, 1 and 2 respectively. Participants chose one of several possible frequencies in response to how often they experience diarrhea, that were recoded as follows: ‘infrequently/never’ = 0, ‘once a week or less’ = 1, ‘more than once a week’ = 2 and ‘daily’ = 2. Similarly, alcohol use (no. of drinks per day) was reported on the following scale which was recoded into corresponding ordinal variables: (0) ‘I do not drink’, (1) ‘1-2 drinks’: (2) ‘3-4 drinks’: (3) ‘5-6 drinks’: (4) ‘More than 6 drinks’. Current tobacco use and prescription medication were both modelled as binary variables (yes/no). Finally, for dietary variables (fruit, vegetables, grains, and sweets intake), participants chose one of the following responses, which were then recoded to the corresponding ordinal variables: (grains): (0) ‘Zero/less than 1 per day’: (1) ‘1-2’: (2) ‘3-4’: (3) ‘5-6’: and (4) ‘7 or more’. (fruits, vegetables): (0) ‘Zero/less than 1 per day’:’(1) ‘1’: (2) ‘2-3’: (3) ‘4-5’: (4) ‘6 or more’. (chocolates/sweets): (0) ‘Less than once per month’: (1) ‘1-3 times per month’: (2) ‘Once per week’: ‘(3) ‘2-4 times per week’: (4) 5-6 times per week’: (5) Once per day’: (6) 2-3 times per day’: (6) ‘4-5 times per day’: (6) ‘6+ times per day’. Sleep was reported as the average amount of sleep you get a day on a three-point scale: (0) ‘Less than 6 hours’: (1) ‘7 to 9 hours’: (2) ‘More than 9 hours’. As the Arivale cohort consists of self-enrolled participants, the response rates for different questionnaires vary. The number of missing values for each response is reported in Table S2.

### Health Measures in the MrOS Cohort

We utilized four different health measures that were collected on MrOS participants during their fourth follow-up visit. Medication use, self-perceived health, and the Life-Space score (LSC) were all self-reported. Self-perceived health captured each individual’s rating of their own health compared to other individuals their own age. The implementation of the LSC in the MrOS cohort has been described in detail previously ^33^. Briefly, LSC can range from 0 (restricted to one’s bedroom) to 120 (traveled outside one’s town daily without assistance). We defined healthy individuals as those in the top tertile of the LSC cohort distribution. This corresponded to an LSC value of ≥96. Walking speed was calculated based on the time it took each participant to walk 6 meters (m/s). Like with the LSC, we defined healthy individuals based on walking speed if their speed was in the top tertile (≥1.17). A total of 7 MrOS participants did not have available walking speed data. This is due to either the participants not coming to the clinic, or not being able to attempt the test. These individuals were classified in the walking speed low group in our analysis.

### Statistical Analysis

Depending on the statistical approach, analysis was conducted using either R (v 3.6) or Python (v 3.7). The relationship between the calculated uniqueness measure and age in the Arivale cohort was modeled using Ordinary Least Square (OLS) linear regression (Python) where square root transformed Bray-Curtis uniqueness was modeled as the dependent variable and each age decade was compared to the youngest reference group (<30 years), adjusting for sex, BMI, and Shannon diversity. We chose to adjust for Shannon diversity because, in our analysis, it was associated with both age and microbiome uniqueness (higher alpha diversity makes you more likely to be unique). We wanted to assess the significance of our dissimilarity pattern independent of changes in alpha diversity seen with age and previously reported in literature. The same adjustment was not made for MrOS participants, since Shannon diversity showed no association with age in that cohort. Pearson/Spearman correlations were also used to assess the strength of the relationship between different measures of ß- and α-diversity and age across all cohorts using the Python statistical functions package (scipy.stats). When assessing the relationship between clinical, lifestyle, and demographic variables with gut microbial uniqueness, Bray-Curtis uniqueness values greater or less than 3 standard deviations from the mean were removed. OLS linear regression was then used to assess the individual relationship between each factor and square root transformed Bray-Curtis gut microbial uniqueness, with microbiome vendor included as a covariate. Percent variance explained by each factor was calculated by taking the percent variance explained by the complete OLS model (variable of interest and microbiome vendor) and subtracting the percent variance explained by microbiome vendor alone. The same analysis was then repeated with age included as a covariate (age-adjusted models). To investigate potential effect modification of sex on the identified gut microbiome aging pattern, an OLS model was generated with a sex*age interaction term predicting square root transformed Bray-Curtis uniqueness, adjusted for sex, age, BMI, microbiome vendor and Shannon diversity. Sex-specific ß-coefficients were estimated by first stratifying the cohort by sex and then fitting OLS models for men and women separately, adjusting for the same covariates as the combined model. Age was scaled and centered prior to this analysis. When investigating the relationship between plasma metabolite concentrations and gut microbial uniqueness, each metabolite was log transformed and subsequently scaled and centered. The square root transformed Bray-Curtis uniqueness score was then regressed against each metabolite individually, adjusting for microbiome vendor, sex, age, age^2^, a sex*age interaction term, BMI, and Shannon diversity using OLS regression. In each instance where multiple hypotheses were tested, type I error was controlled for using the Bonferroni method (P<0.05). In the MrOS Cohort, correlation between Bray-Curtis Uniqueness and age was calculated using the Python statistical functions package (scipy.stats) using the square root transformed uniqueness score. Mortality analysis was conducted in R using the package survival (v 2.44-1.1). Relative *Bacteroides* abundance (after rarefaction) and uniqueness scores were scaled and centered prior to survival analysis. Cox-proportional hazard regression models were generated assessing the relationship between survival and Relative *Bacteroides* abundance or Bray-Curtis uniqueness independently, adjusting for clinical site, batch (discovery/validation) and age, and adjusting for clinical site, age, BMI, self-perceived health (excellent, good, <good), diagnosis of congestive heart failure, and batch in which stool samples were processed (discovery/validation).

## Supplementary Information

**Fig. S1.**
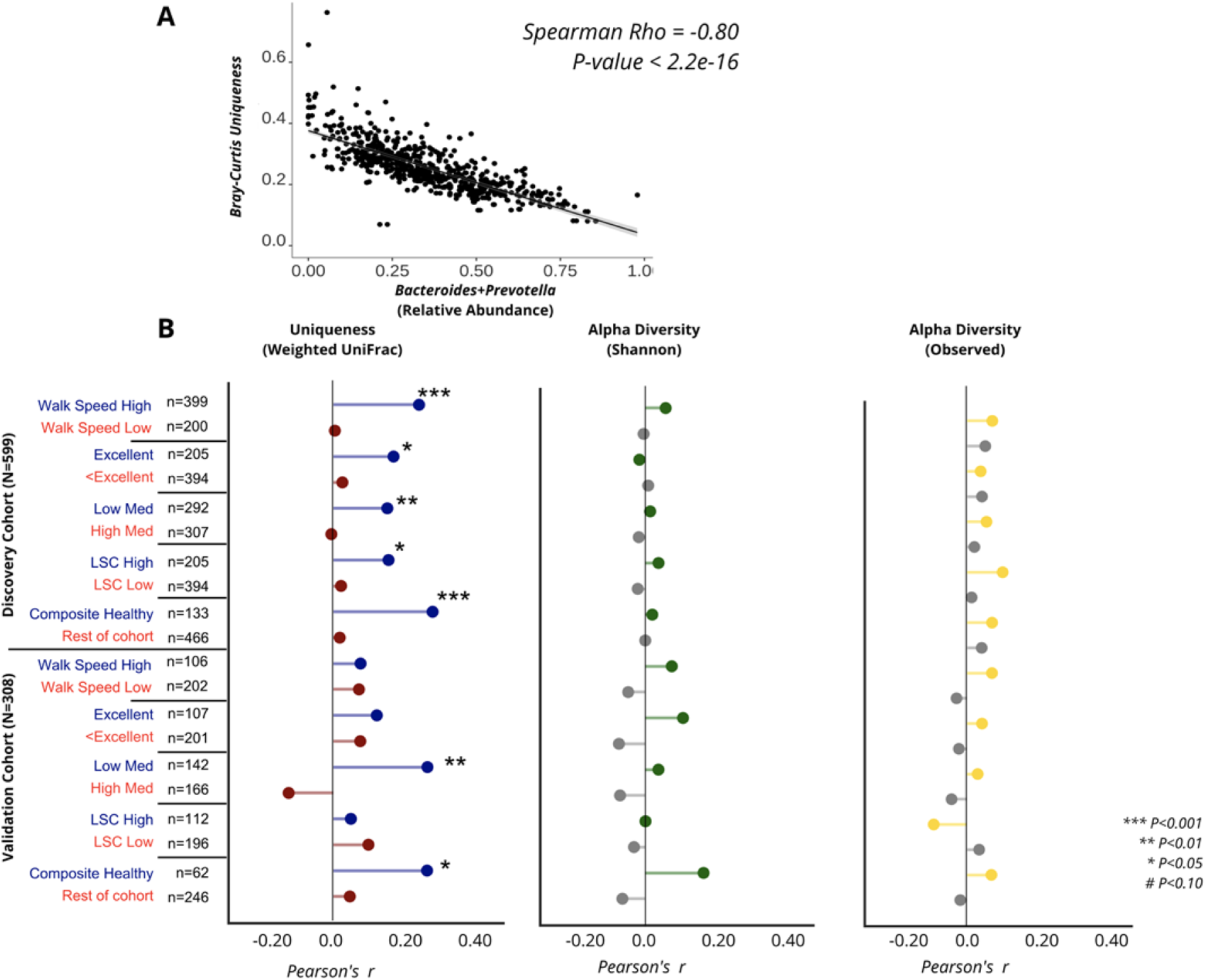
Associations between age and gut microbiome measures across health stratifications in the MrOS cohort. **(A)** Scatter plot demonstrating the negative association of the relative abundance of the sum of *Bacteroides*+*Prevotella* and gut microbial uniqueness. The line shown is the y∼x regression line, while the shaded region corresponds to the 95% confidence intervals for the slope of the line. **(B)** Plots demonstrating the strength of correlation between age and gut microbiome measures. The blue/red panel corresponds to the calculated Weighted UniFrac (ß-diversity) uniqueness score, while the grey/green and grey/yellow panels correspond to Shannon diversity and Observed species (α-diversity measures), respectively. Significant correlations are indicated with pound signs and asterisks.

**Table S1.** Arivale cohort characteristics stratified by microbiome vendor. Statistical tests used to compare groups are as follows: independent samples t-tests were used for comparing age, body mass index (BMI), high density lipoprotein (HDL), low density lipoprotein (LDL), blood triglycerides and Bray-Curtis Uniqueness; nonparametric Mann–Whitney U tests were used to compare Shannon diversity and Observed OTUs; χ2 tests were used to compare sex (percentage female) and race (percentage non-white). *P-Values* <0.05 are colored in red.

|  | Vendor A (N=2539) | Vendor B (N=1114) | P-Value |
| --- | --- | --- | --- |
| Mean age (range) | 48.4 (18-87) | 48.3 (19-82) | NS |
| Mean BMI (s.d.) | 27.1 (5.9) | 27.3 (6.1) | NS |
| Sex (% female) | 58.60% | 60.90% | NS |
| Non-white (%) | 20.40% | 22.00% | NS |
| Mean HDL (mg dl<sup>-1</sup>) (s.d.) | 61.8 (18.8) | 61.7 (18.4) | NS |
| Mean LDL (mg dl<sup>-1</sup>) (s.d.) | 114.0 (33.4) | 114.0 (34.7) | NS |
| Mean blood triglycerides (mg dl<sup>-1</sup>) (s.d.) | 104.9 (59.6) | 105.3 (58.0) | NS |
| Median Shannon diversity [IQR] | 4.34 [4.04-4.60] | 4.31 [3.99-4.57] | 0.032 |
| Median Observed OTUs [IQR] | 276 [217-247] | 287.5 [226-371] | 4.60E-05 |
| Mean Bray-Curtis uniqueness (s.d.) | 0.24 (0.06) | 0.26 (0.07) | 3.94E-08 |

**Table S2.**
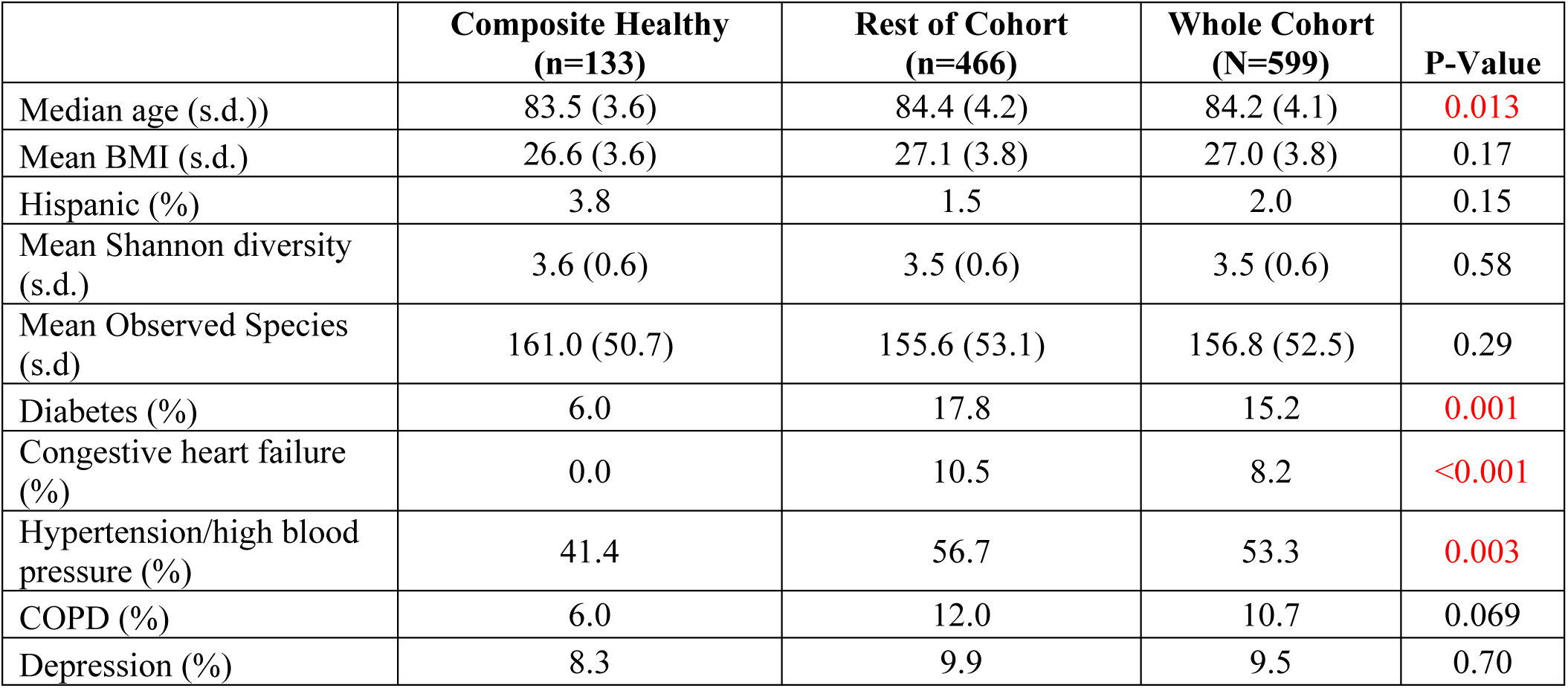
MrOS discovery cohort characteristics stratified into composite healthy and remainder of cohort. Statistical tests used to compare groups are as follows: independent samples t-tests were used for comparing age, body mass index (BMI), Shannon diversity and Observed Species; χ^2^ or Fisher’s exact (if assumptions of χ^2^ were not met) tests were used to compare ethnicity (percentage Hispanic), and prevalence of each of the specified diseases. P-values <0.05 are colored in red.

**Table S3.** Associations between Bray-Curtis gut microbiome uniqueness and clinical, demographic, and diet/lifestyle/health measures. ‘P-Value’ corresponds to the unadjusted P-Value of the ß-coefficient (B-Coefficient column) for each analyte from an OLS model adjusted for gut microbiome vendor. ‘r_squared’ reflects the percent of variance explained beyond microbiome vendor for each analyte independently. ‘Missing’ shows the number of missing observations for each analyte. ‘Age adj. B-coefficient’ and ‘Age adj. P-value’ correspond to the ß-coefficient and P-Value for each analyte adjusting for gut microbiome vendor and age. Values highlighted in red are statistically significant after multiple-hypothesis correction (Bonferroni P-Value<0.05).

| Analyte | P-Value | r_squared | B-coefficient | Missing | Age adj. B-coefficient | Age adj. P-Value |
| --- | --- | --- | --- | --- | --- | --- |
| Age | 5.89E-33 | 3.85901529 | 0.00093261 | 0 | NA | NA |
| Prescription Med | 0.00017481 | 1.53219831 | 0.01510618 | 914 | 0.00341449 | 0.08545254 |
| HDL | 3.03E-08 | 0.8659935 | 0.00549509 | 104 | 0.003964398 | 5.47E-05 |
| n6/n3 | 3.66E-08 | 0.855769 | -0.0054647 | 104 | -0.003148386 | 0.00156995 |
| Vitamin D | 1.08E-07 | 0.79702949 | 0.00527528 | 104 | 0.003159677 | 0.00144512 |
| Alcohol | 1.36E-05 | 0.5608151 | -0.0070819 | 3349 | -0.002982965 | 0.0028601 |
| Homocysteine | 0.00014395 | 0.4087152 | 0.00377064 | 104 | 0.001589201 | 0.10941824 |
| BMI | 0.00044766 | 0.36457364 | -0.0005975 | 260 | -0.003869322 | 0.00010072 |
| Triglycerides | 0.00077075 | 0.32005925 | -0.0033549 | 104 | -0.004164096 | 1.97E-05 |
| Diarrhea | 0.00238587 | 0.26867461 | -0.0040083 | 3415 | -0.001701617 | 0.08690565 |
| HbA1c | 0.00715616 | 0.204807 | 0.00272545 | 104 | 6.97E-05 | 0.94455438 |
| Race(ref.white) | 0.01700183 | 0.16061072 | 0.0058388 | 88 | 0.000687553 | 0.48477659 |
| Antibiotics | 0.05402505 | 0.14773123 | 0.00642737 | 2500 | 0.001844306 | 0.10335017 |
| Sex | 0.02355609 | 0.14101656 | 0.00452064 | 0 | 0.00236889 | 0.01379305 |
| Sweets | 0.27161453 | 0.13414917 | -0.0012896 | 902 | -4.21E-05 | 0.98258201 |
| ALAT | 0.04087102 | 0.11844347 | -0.0020499 | 104 | -0.002165898 | 0.02620547 |
| CRP | 0.04604621 | 0.11274027 | -0.0019787 | 104 | -0.001966477 | 0.04354322 |
| LDL | 0.04967172 | 0.10913342 | -0.0019505 | 104 | -0.003130326 | 0.00137829 |
| Glucose | 0.10416215 | 0.07481646 | 0.00163161 | 104 | -0.000440919 | 0.6562693 |
| Tobacco | 0.11771405 | 0.07460265 | -0.0077207 | 3264 | -0.00115745 | 0.2504751 |
| Globulin | 0.13060992 | 0.06475052 | -0.0015033 | 104 | -0.000324745 | 0.74022842 |
| ALP | 0.18418122 | 0.0499728 | 0.00133152 | 104 | -0.000589978 | 0.55024706 |
| GGT | 0.25345563 | 0.03695739 | -0.0011321 | 104 | -0.0023493 | 0.01643053 |
| Grains | 0.32021289 | 0.029105 | -0.0012634 | 3379 | -0.000165682 | 0.86721005 |
| Fruits | 0.34161948 | 0.02630667 | 0.00113363 | 3422 | -0.000172467 | 0.86142995 |
| Insulin | 0.48819587 | 0.0136181 | -0.0006939 | 104 | -0.001210903 | 0.21445978 |
| Vegetables | 0.84610634 | 0.0010959 | 0.00022377 | 3422 | -0.000193127 | 0.84450268 |
| Sodium | 0.89643796 | 0.0004802 | 0.00012949 | 104 | -0.000934367 | 0.33958969 |
| Sleep | 0.93138998 | 0.000316 | 0.00021369 | 2335 | 0.000427359 | 0.71350207 |
| HOMA-IR | 0.95318368 | 9.77E-05 | -5.85E-05 | 104 | -0.000840639 | 0.38949581 |
| Creatinine | 0.98900622 | 5.38E-06 | -1.37E-05 | 104 | -0.000570523 | 0.55888158 |

**Table S4.** Associations between Bray-Curtis gut microbiome uniqueness and plasma metabolites. Only metabolites with P-value<0.01 are shown. **‘**P-Value’ corresponds to the covariate adjusted P-value of the ß-coefficient (Adj. B-coefficient). ‘Corrected P-Value’ corresponds to the Bonferroni corrected P-values. Metabolites significantly associated with Bray-Curtis uniqueness after adjusting for covariates and multiple hypothesis correction are highlighted in red. * indicates metabolites that were confidently identified on the basis of mass spectrometry data, but for which no reference standards are available to verify the identity.

| Metabolite | P-Value | Corrected P-Value | Adj. B-coefficient |
| --- | --- | --- | --- |
| phenylacetylglutamine | 3.65E-20 | 2.38E-17 | 0.015774767 |
| p-cresol glucuronide* | 6.76E-13 | 4.40E-10 | 0.011978401 |
| 6-hydroxyindole sulfate | 7.06E-09 | 4.60E-06 | 0.009597508 |
| 3-indoxyl sulfate | 5.24E-07 | 0.00034152 | 0.008269926 |
| lithocholate sulfate | 7.39E-07 | 0.00048191 | 0.008153816 |
| p-cresol sulfate | 5.06E-06 | 0.00329597 | 0.007949108 |
| indoleacetate | 9.55E-06 | 0.00622582 | 0.007363627 |
| taurolithocholate 3-sulfate | 6.67E-05 | 0.04347711 | 0.006750892 |
| trimethylamine N-oxide | 8.02E-05 | 0.0522833 | 0.006676801 |
| glycodeoxycholate 3-sulfate | 0.00011043 | 0.07199921 | 0.006384711 |
| 1,5-anhydroglucitol (1,5-AG) | 0.00019356 | 0.12620424 | -0.006192382 |
| biliverdin | 0.00026726 | 0.17425604 | -0.006295182 |
| carotene diol (1) | 0.00048694 | 0.3174862 | -0.006178362 |
| 4-ethylcatechol sulfate | 0.00059272 | 0.38645107 | 0.005838135 |
| threonate | 0.00095377 | 0.6218559 | -0.005560642 |
| dodecanedioate (C12-DC) | 0.00121002 | 0.78893024 | 0.005391288 |
| N-acetylputrescine | 0.00448236 | 1 | 0.004753617 |
| carotene diol (3) | 0.00272636 | 1 | -0.005152625 |
| ergothioneine | 0.00708238 | 1 | -0.004510327 |
| androstenediol (3alpha, 17alpha) monosulfate (2) | 0.00935419 | 1 | -0.004964594 |
| 3-hydroxy-2-ethylpropionate | 0.00904741 | 1 | 0.004420961 |
| 4-ethylphenylsulfate | 0.00586104 | 1 | 0.004664238 |
| propionylglycine | 0.00191267 | 1 | 0.005169723 |
| isobutyrylcarnitine (C4) | 0.00383686 | 1 | 0.004969195 |
| tartronate (hydroxymalonate) | 0.00388 | 1 | -0.004950944 |
| cys-gly, oxidized | 0.00874903 | 1 | 0.004322096 |
| 4-acetamidobutanoate | 0.0028892 | 1 | 0.00512429 |
| bilirubin (Z,Z) | 0.00233088 | 1 | -0.005203709 |
| cortisol | 0.00512227 | 1 | 0.004670237 |
| 5-methylthioadenosine (MTA) | 0.00432121 | 1 | 0.00487853 |
| androstenediol (3beta,17beta) disulfate (2) | 0.0078715 | 1 | -0.005096967 |
| 2,3-dihydroxy-5-methylthio-4-pentenoate (DMTPA)* | 0.00233228 | 1 | 0.005882396 |

**Table S5.**
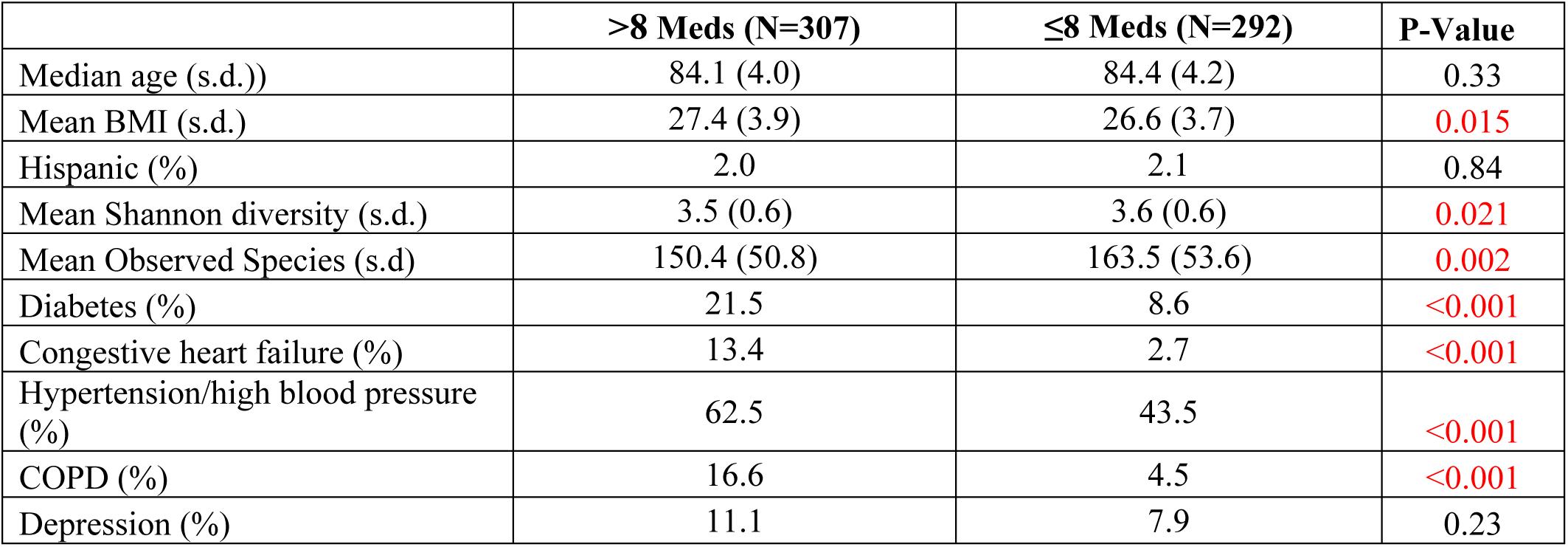
MrOS discovery cohort characteristics stratified by medication use. Statistical tests used to compare groups are as follows: independent samples t-tests were used for comparing age, body mass index (BMI), Shannon diversity and Observed species; χ^2^ or Fisher’s exact (if assumptions of χ^2^ were not met) tests were used to compare ethnicity (percentage hispanic) and prevalence of each of the specified diseases. *P-Values* <0.05 are colored in red.

|  | >8 Meds (N=307) | ≤8 Meds (N=292) | P-Value |
| --- | --- | --- | --- |
| Median age (s.d.)) | 84.1 (4.0) | 84.4 (4.2) | 0.33 |
| Mean BMI (s.d.) | 27.4 (3.9) | 26.6 (3.7) | 0.015 |
| Hispanic (%) | 2.0 | 2.1 | 0.84 |
| Mean Shannon diversity (s.d.) | 3.5 (0.6) | 3.6 (0.6) | 0.021 |
| Mean Observed Species (s.d) | 150.4 (50.8) | 163.5 (53.6) | 0.002 |
| Diabetes (%) | 21.5 | 8.6 | <0.001 |
| Congestive heart failure (%) | 13.4 | 2.7 | <0.001 |
| Hypertension/high blood pressure (%) | 62.5 | 43.5 | <0.001 |
| COPD (%) | 16.6 | 4.5 | <0.001 |
| Depression (%) | 11.1 | 7.9 | 0.23 |

